# Inhibition of norepinephrine signaling during a sensitive period disrupts locus coeruleus circuitry and emotional behaviors in adulthood

**DOI:** 10.1101/287243

**Authors:** Qingyuan Meng, Alvaro L. Garcia-Garcia, Alex Dranovsky, E. David Leonardo

## Abstract

Deficits in arousal and stress responsiveness span numerous psychiatric developmental disorders including depression and anxiety. Arousal is supported by norepinephrine (NE) released from locus coeruleus (LC) neurons onto cortical and limbic areas. During development, the NE system matures in concert with increased exploration of the animal’s environment. While several psychiatric medications target the LC-NE system, the possibility that its modulation during discreet developmental periods can have long-lasting consequences for mental health has not been explored. We used a pharmacogenetic strategy in mice to reversibly inhibit NE signaling during brief developmental periods to determine the long-lasting impact on adult circuits mediating emotional behavior. We also examine whether disruption of NE signaling during development results in permanent changes within the adult LC-NE system. Finally, we test whether developmental exposure to the α-2 receptor agonist guanfacine recapitulates the effect seen with our pharmacogenetic strategy. Our results reveal a sensitive period (postnatal days 10-21) during which alterations in NE signaling result in long-term changes in adult emotional behavior. Changes in NE signaling during this sensitive period results in changes in stress-related LC neuron activity, alterations in α-2 autoreceptor function, and circuit-specific molecular changes in LC-NE target regions in adulthood. Treating animals with guanfacine during the sensitive period produced similar results. Our findings indicate an early critical role for NE in sculpting brain circuits that support adult emotional function.

## Introduction

The locus coeruleus (LC), a small densely packed brain stem nucleus, is the primary source of norepinephrine (NE) in the brain (Sara, 2009). The LC-NE system is involved in a wide range of normal behavioral and physiological responses such as arousal (Sara, 2009; Carter et al., 2010; Snyder et al., 2012) and is a major component of the centrally mediated stress response (Valentino and Van Bockstaele, 2008; Olson et al., 2011). Behaviors supported by normal LC-NE function are disrupted numerous psychiatric disorders (Schwarz and Luo, 2015). Indeed, dysregulation within the NE system has been implicated in the pathophysiology and treatment of anxiety and depression (Goddard et al., 2010). For example, α-2 receptor agonists and antagonists alleviate or exacerbate anxiety symptoms respectively (Charney, 2003). Moreover, fewer NE transporters (NET) have been reported in the LC of post-mortem brain tissue obtained from depressed patients (Klimek et al., 1997; Cottingham and Wang, 2012). Together, this evidence implicates LC dysfunction in the biology of neuropsychiatric conditions.

The ability of NE to modulate brain functions develops postnatally as the maturing LC begins to exhibit activity in response to environmental changes (Nakamura et al., 1987). For example, LC-NE signaling is critical for early life sensory and odor-based attachment learning (Rangel and Leon, 1995; Raineki et al., 2010). These processes are complete by postnatal day 10-15 (P10-P15), just as LC autoinhibition through α-2A autoreceptors comes online (Murrin et al., 2007; Raineki et al., 2010). Interestingly, in rodents, the time period that follows (P15-P21) coincides with the emergence of normal exploration and habituation (Ba and Seri, 1995). These and other milestones of LC function occur in the context of steadily increasing NE from birth until P30-40 when adult concentrations are reached (Loizou and Salt, 1970; Konkol et al., 1978; Morris et al., 1980). Genetic models also support the notion that disrupting NE signaling has developmentally distinct consequences. While α-2A receptor KO mice display an anxiety and depression-like phenotype (Schramm et al., 2001; Lahdesmaki et al., 2002), adult suppression of LC α-2A autoreceptors leads to the opposite outcome (Shishkina et al., 2002). Moreover, drugs that target the NET for the treatment of depression are ineffective in juveniles but highly effective in adults (Geller et al., 1999; Hazell et al., 2000), suggesting that the effects of modulating NE signaling on emotion related circuits differ across the lifespan. Thus, given that NE has distinct roles in circuit maturation, we hypothesized that briefly disrupting NE function during different developmental periods would have distinct consequences.

Here, we tested whether disruption of normal NE signaling during brief developmental time periods has long-term consequences for normal adult NE function in the brain. We found that pharmacogenetic inhibition of NE signaling during P10-P21, but not earlier or later resulted in molecular adaptations within the LC-NE system along with a depression-like phenotype with some anxiety features. Administration of the α-2A agonist guanfacine, a medication commonly used in children, during this sensitive period phenocopied the pharmacogenetic results highlighting the potential vulnerability of developing brain systems to psychoactive medications.

## Materials and Methods

Detailed information on experimental procedures is provided in Supplementary Information.

### Generation of DBH-hM4Di mice

RC∷PDi (Cre-dependent inhibitory DREADD [hM4Di] receptor) mice have been described (Ray et al., 2011) and were a generous gift from Susan Dymecki. Tg(Dbh-cre)KH212Gsat/Mmcd (DBH-Cre) mice, identification number 032081-UCD, were obtained from the Mutant Mouse Regional Resource Center, a NCRR-NIH funded strain repository, and was donated to the MMRRC by the NINDS funded GENSAT BAC transgenic project. RC∷PDi and DBH-Cre lines were crossed to generate the DBH-Cre-RC∷PDi line (DBH-hM4Di+), which was maintained on a mixed C57BL/6 and 129S6/Sv background.

### Drug treatment and administration

#### Clozapine-N-oxide (CNO)

CNO was obtained from the NIH as part of the Rapid Access to Investigative Drug Program funded by the NINDS. For adult acute administration, CNO was injected at a dose of 5 mg/kg in 1% DMSO in 0.9% saline intraperitoneally (i.p.).

For developmental interventions, DBH-hM4Di^+^ male mice were treated daily with vehicle or CNO i.p. (5 mg/kg in 1% DMSO and 0.9% saline) during different time windows (P2–P9/ P10-P21/ P33-P45/ P56-P67). Specifically, 10 μL per 1 gr of mouse body weight was injected from a 0.5 mg/mL stock solution. For pre-weaning treatments (P21), the entire litters were removed from dams and placed in a small tray containing bedding from the respective home cage. The tray was placed on a scale allowing us to measure the weight of an individual pup when removing it for injection. Mice were injected in a random order and immediately placed back in the home cage. All mice within a litter were assigned to the same treatment.

#### Systemically induced adult hypothermia

clonidine (0.5 mg/kg in 0.9% saline) and 8-OH-DPAT (1 mg/kg in 0.9% saline) (Sigma–Aldrich, St. Louis, MO) (Richardson-Jones et al., 2011) were administered i.p.

#### Guanfacine

1 mg/kg/day in 0.9% saline was administered i.p.

### Behavioral and physiological studies

13–15-week-old male mice were tested over 4–5 weeks. Anxiety tests were completed before other behavioral tests.

### Open-field test

Exploration of a novel open field was measured for 30 min as previously described (Garcia-Garcia et al., 2015; Garcia-Garcia et al., 2017). The center of the arena was defined as a square area occupying the center 50% of the total arena. Dependent measures were total path length (cm) and percent distance in the center (distance travelled in the center divided by the total distance travelled).

### Elevated-plus maze

Animals were placed into the central area facing one open arm and allowed to explore the maze for 5 min as previously described (Garcia-Garcia et al., 2015; Garcia-Garcia et al., 2017). Illumination of 90–100 lux was used. The videos were analyzed with TopScan software (Clever Sys Inc, Reston, VA). Dependent measures were time in the open arms and percent time in the open arms (time in the open arms divided by the total time).

### Sucrose Preference

An 8-day sucrose preference protocol was performed as previously described (Garcia-Garcia et al., 2015; Garcia-Garcia et al., 2017). On days 1 and 2, mice were presented with two water filled bottles (water/water) for 2 h and 1 h respectively. On days 3 and 4, both bottles contained 1% sucrose in water (sucrose/sucrose) for 1 h and 30 min respectively. On choice days 5–8 (testing days), one bottle contained water and the other 1% sucrose solution for 30 min each day (water/sucrose). Daily preference was calculated as: ((weight bottle 1)/ (weight bottles (1 + 2)) ×100).

### Forced-swim test

Behavioral response to forced swimming was assayed as described previously (Garcia-Garcia et al., 2015; Garcia-Garcia et al., 2017). Mice were placed into clear plastic buckets 20 cm in diameter and 23 cm deep filled 2/3 of the way with 26°C water and videotaped from the side for 6 min. Only the last 4 minutes were scored. Scoring was done using an automated Viewpoint Videotrack software package (Montreal, Canada), which was validated before by manual scoring. Dependent variable was immobility.

### Hypothermia

Mice were singly housed in clean cages for an hour and three baseline temperatures were taken (-60 min, -30 min, 0min). Immediately after the last baseline measurement (0 min), animals received CNO or clonidine as described above. Body temperature was assessed rectally using a lubricated probe (Thermalert TH-5 thermal monitor, Physitemp, Clifton, NJ) every 10 min for 60 or 120 minutes.

### Corticosterone measurements

Blood was collected during the dark-light transition and 12 hours later during the light-dark transition as previously described (Garcia-Garcia et al., 2015; Garcia-Garcia et al., 2017). For stress-evoked corticosterone, experiments were performed starting at 12.00. Mice were exposed to a forced swim stressor for 6 min and blood was drawn from the submandibular vein 12 min later. Blood was centrifuged and plasma was isolated and stored until processed. Corticosterone levels were assessed by ELISA from plasma samples (Enzo Life Sciences, Farmingdale, NY) (Garcia-Garcia et al., 2015; Garcia-Garcia et al., 2017).

### C-fos Immunohistochemistry

C-fos was induced as previously described by a forced swim stressor (Garcia-Garcia et al., 2017). After tissue processing, LC sections were stained using rabbit anti-c-fos antibody (1:5000, Millipore) and sheep anti-tyrosine hydroxylase (TH) (1:1000, Abcam) as previously described (Garcia-Garcia et al., 2017).

### Quantitative PCR

Total RNA was extracted using TRIzol and SuperScript® III First-Strand Synthesis System was used to synthesize cDNA, and PCR was performed and quantified using SYBR Green real-time PCR Master Mix (Life Technologies, Grand Island, NY). Primers used in the real-time qPCR are in the supplemental information.

### Brain neurotransmitter levels

NE concentrations were determined by HPLC as previously described (Garcia-Garcia et al., 2015). Specifically, all assays were carried out on a Waters Xevo TQ MS ACQUITY UPLC system (Waters, Milford, MA, USA). Concentrations of compounds in the samples were quantified by comparing integrated peak areas against those of known amounts of purified standards. Loss during extraction was accounted for by adjusting for the recovery of the internal standard added before extraction. The results were normalized by sample weight.

### Statistical Analysis

All statistical analyses were performed using Stat View (SAS Institute Inc.). Final group numbers are shown in the figure legends. The results were expressed as mean ± SEM. *p*<0.05 was used as the threshold for significance. Group differences were analyzed using a one-way analysis of variance (ANOVA) unless otherwise stated. Repeated measures ANOVA was used for hypothermia and sucrose preference experiments.

## Results

### Manipulation of NE Neuronal Activity via Conditional Expression of DREADDs in Mice

To interfere with NE signaling *in vivo*, we crossed RC∷PDi mice (Ray et al., 2011) to a dopamine β-hydroxylase cre line (DBH-Cre) (Gong et al., 2007). The resulting DBH-cre;RC:PDi (DBH-hM4Di^+^) mice express hemaglutinin (HA) tagged hM4Di (inhibitory DREADD) in NE neurons (Figure 1A, B). LC immunostaining against Tyrosine hydroxylase (TH) and HA confirmed selective DREADD expression in virtually all LC-NE neurons (Figure 1B). This approach allows for targeted and time limited inhibition of NE neurons by activation of hM4Di with the biologically inert synthetic ligand CNO (Armbruster et al., 2007).

**Figure 1.**
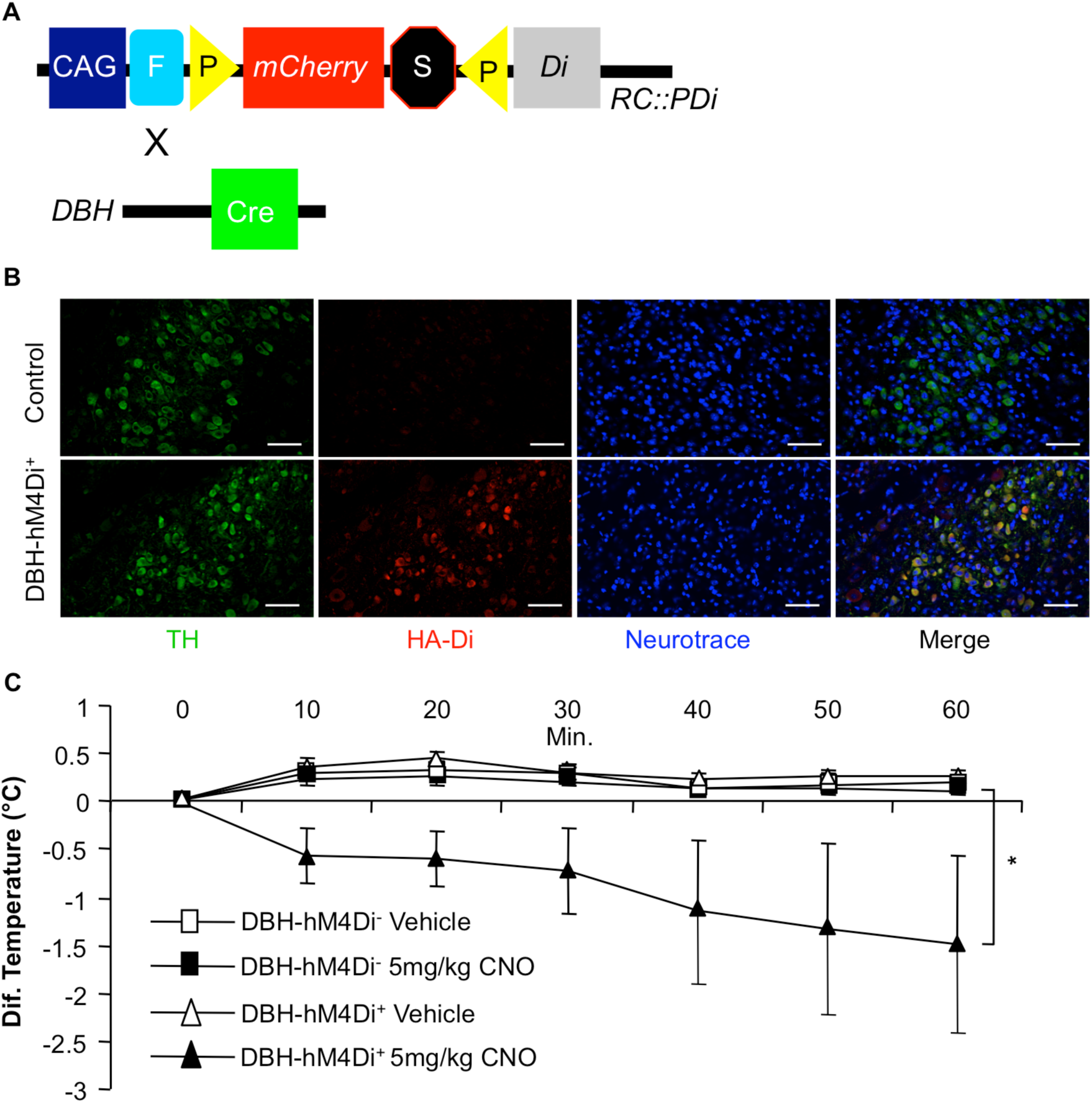
CNO interferes with LC function in DBH-hM4Di mice. **(A)** The inhibitory DREADD hM4Di (Di) was expressed in NE neurons by crossing the RC∷PDi (hM4Di) line with a dopamine β-hydroxylase cre line (DBH-Cre), resulting in a Rc:PDi;DBH-cre (DBH-hM4Di^+^) line that expresses hM4Di in NE cells. **(B)** Immunostaining for TH, HA-tag, and Neurotrace in the LC of DBH-hM4Di^+^ animals confirm the specificity of hm4Di expression in NE neurons. Scale bars=100 µm. (**C)** Hypothermic response to an injection of vehicle or 5 mg/kg of CNO in DBH-hM4Di^−^ and DBH-hM4Di^+^ mice in adulthood (two-way ANOVA repeated measures genotype x treatment interaction: F_3,23_=4.364, p<0.05; n=3/group). Time course is shown in minutes. Temperature is shown as change from baseline. Means are represented as ±SEM. (*p<0.05).

We next took advantage of the fact that α2-A receptor agonists cause a modest hypothermia in adult animals by inhibiting NE neurons (Madden et al., 2013). We thus tested whether CNO injection would also cause hypothermia by inhibiting NE neurons. Indeed, administration of CNO (5 mg/kg) but not vehicle resulted in sustained hypothermia in adult DBH-hM4Di^+^ mice. No effect was seen in DBH-hM4Di^−^ mice treated with vehicle or CNO (Figure 1C). This result demonstrates that hM4Di receptors expressed in NE neurons are functional.

### Repeated pharmacogenetic inhibition of NE neurons between P10 and P21 results in increased anxiety in the open-field and depression-like behavioral phenotype later in life

We next hypothesized that normal NE function in early development is critical to establish later behavioral responses. We tested this by disrupting NE function through daily administration of CNO (5 mg/kg) to DBH-hM4Di^+^ mice during three discrete postnatal developmental periods (Figure 2A). The LC exhibits bursts of activity in response to external stimulation by the first week of life (Nakamura et al., 1987). Between P10 and weaning (P21), robust spontaneous LC firing emerges and α-2A receptor negative feedback is established (Happe et al., 2004). We therefore selected P2-P9, P10-21, and an adult reference period (P56-P67) for our experimental manipulations.

**Figure 2.**
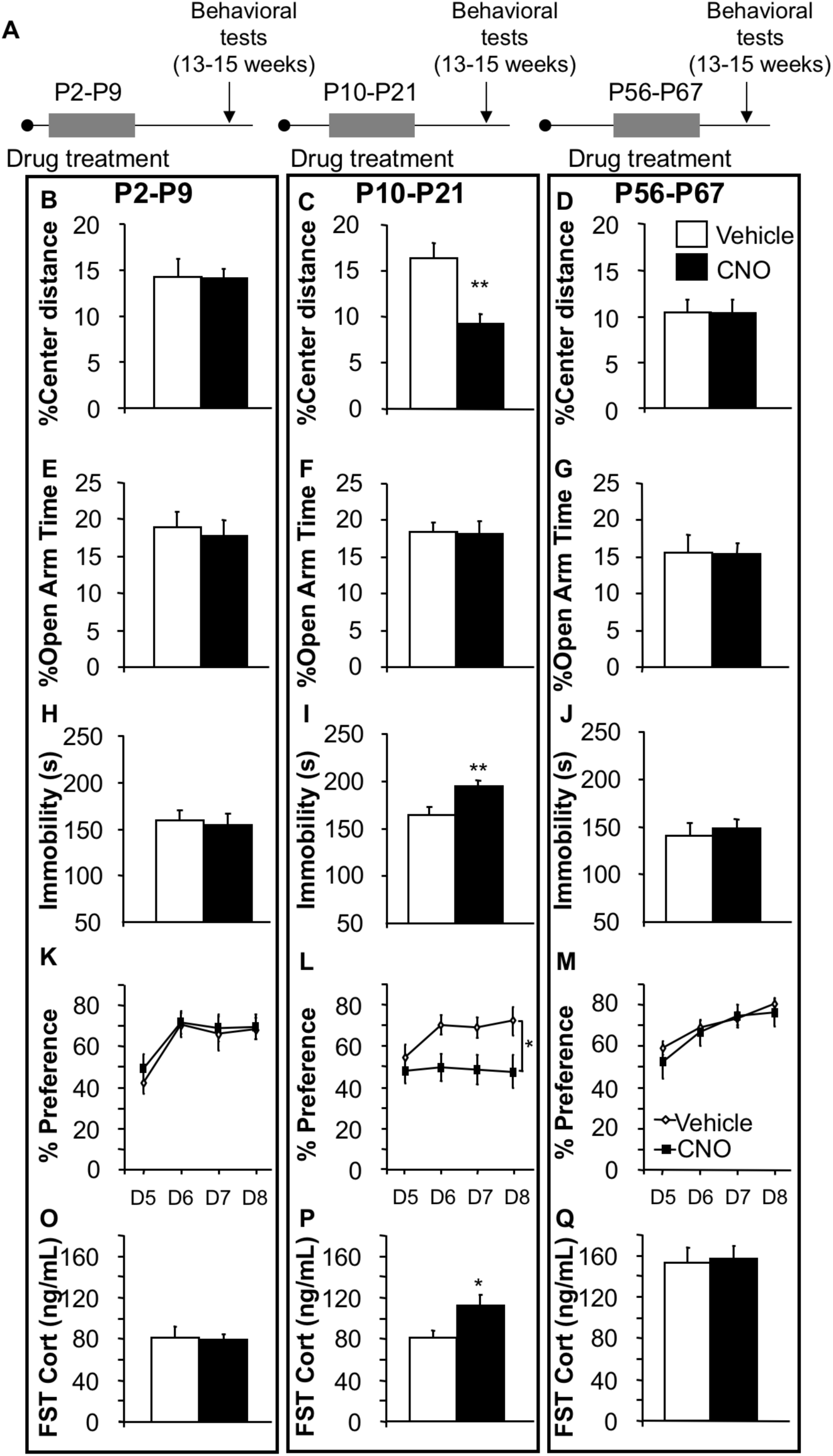
Enhanced anxiety and depression-like behavior in adult mice after pharmacogenetic inhibition of NE neurons between P10-P21 but not after P2-P9 or P56-P67. **(A)** Experimental timeline. (**B-D)** Decreased percent center distance in the open-field after P10-P21 intervention but not after P2-P9 or P56-P67 (one-way ANOVA for main effect of treatment P2-P9: F_1,39_=0.004, p=0.9488; P10-P21: F_1,36_=14.211, p<0.01; P56-P67: F_1,31_=0.002, p=0.9665; n=16-22/group). **(E-G)** No differences were detected in time spent in the open arms in the elevated-plus maze (one-way ANOVA for main effect of treatment P2-P9: F_1,39_=0.140, p=0.7103; P10-P21: F_1,36_=0.001, p=0.981; P56-P67: F_1,31_=0.001, p=0.9714; n=16-22/group). **(H-J)** Immobility in the FST. Differences were observed in the P10-P21 but not other groups when comparing CNO to vehicle mice (one-way ANOVA for main effect of treatment P2-P9: F_1,39_=0.097, p=0.7574; P10-P21: F_1,36_=9.342, p<0.01; P56-P67 F_1,31_=0.186, p=0.6692; n=16-22/group). **(K-M)** Sucrose Preference test. Differences in preference were observed in the P10-P21 group but not others when comparing CNO to vehicle treatment during testing days (Day 5-8) (one-way ANOVA repeated measures for main effect of treatment: P2-P9: F_1,32_=0.735, p=0.3977; P10-P21: F_1,32_=7.256, p<0.05; P56-P67: F_1,26_=0.681, p=0.4168; n=14-22/group). **(O-Q)** Forced swim-stress induced CORT. CORT was significantly increased only in P10-P21 CNO treated group when compared to vehicle treated mice (one-way ANOVA for main effect of treatment: P2-P9: F_1,16_=0.055, p=0.817; P10-P21: F_1,18_=5.051, p<0.05; P56-P67: F_1,18_=0.045, p=0.8341; n=9-11/group). Means are represented as ±SEM. (*p<0.05; **p<0.01). See also Supplementary Figure 1 and 2.

To assess the effects of CNO treatment on exploration and anxiety later in life, we first tested the treated animals at 13-15 weeks of age in the open-field test. Adult DBH-hM4Di^+^ mice treated with CNO during P10-P21 but not during the other time periods, displayed decreased percent distance traveled in the center (Figure 2B-D) with no significant changes in total path traveled (Supplementary Figure 1A-C). In the elevated-plus maze no significant differences were detected between groups in percent time spent in the open arms (Figure 2E-G).

Next, we examined the effect of developmental suppression of NE neurons on the behavior of adult mice in the forced-swim test (FST). Of the three time periods tested, only DBH-hM4Di^+^ mice treated with CNO during P10-P21 displayed increased immobility (Figure 2H-J). We then tested the mice in the sucrose preference test, which measures anhedonia, a core feature of depression (Treadway and Zald, 2011). Once again, only the P10-P21 cohort treated with CNO exhibited reduced sucrose preference during the choice days (Days 5-8) (Figure 2K-M). These results indicate that inhibition of NE neurons during P10-P21, but not during the other time windows, results in increased anxiety in the open-field test along with a depression-like phenotype in the FST and sucrose preference.

Given the role of the NE system in mediating stress responses and in depression, we decided to test the hypothalamic-pituitary-adrenal axis reactivity in the three groups of mice. We used a forced swim stressor to elicit corticosterone (CORT) responses and collected blood samples shortly after. Only the P10-P21 CNO treated group displayed exaggerated stress-induced CORT (Figure 2O-Q) from its respective controls. Importantly, no differences in CORT levels were detected during the diurnal cycle (Supplementary Figure 1D-F).

While robust spontaneous LC activity emerges during the later pre-weaning period, adult NE levels are not reached until adolescence, during the fifth week of life (Murrin et al., 2007). We therefore asked whether animals that underwent NE neuron inhibition during adolescence (P33-P45) would more closely resemble adult- or the juvenile-treated animals with regards to behavioral and physiological consequences. However, CNO treatment during P33-P45 revealed no change in anxiety-like behaviors, depression-related behaviors or stress-induced CORT (Supplementary Figure 2A-H).

### Transient inhibition of NE neurons between P10-P21 has long lasting effects on stress-induced LC-NE neuronal activation

Having observed that interfering with normal NE signaling in the P10-P21 time period results in sustained changes in behavioral and physiological reactivity, we tested whether we could detect sustained changes in the response properties of the LC-NE neurons. To do so, we evaluated c-fos expression in the LC of adult animals in response to a forced-swim stressor. We detected fewer c-fos positive cells amongst NE (TH+) cells in the LC of mice treated with CNO during P10-P21 but not during other time windows (P2-P9, P56-P67) when compared to their respective controls (Figure 3A-C and Supplementary Figure 3B, E, H). No difference in the number of c-fos positive cells in TH negative cells was detected in any of the groups (Supplementary Figure 3C, F, I). Further, the number of TH positive neurons remained unchanged (Supplementary Figure 3A, D, G). Thus, NE signaling inhibition during P10-P21 leads to a decreased stress-induced activation of LC-NE cells in adulthood, without impacting their number.

**Figure 3.**
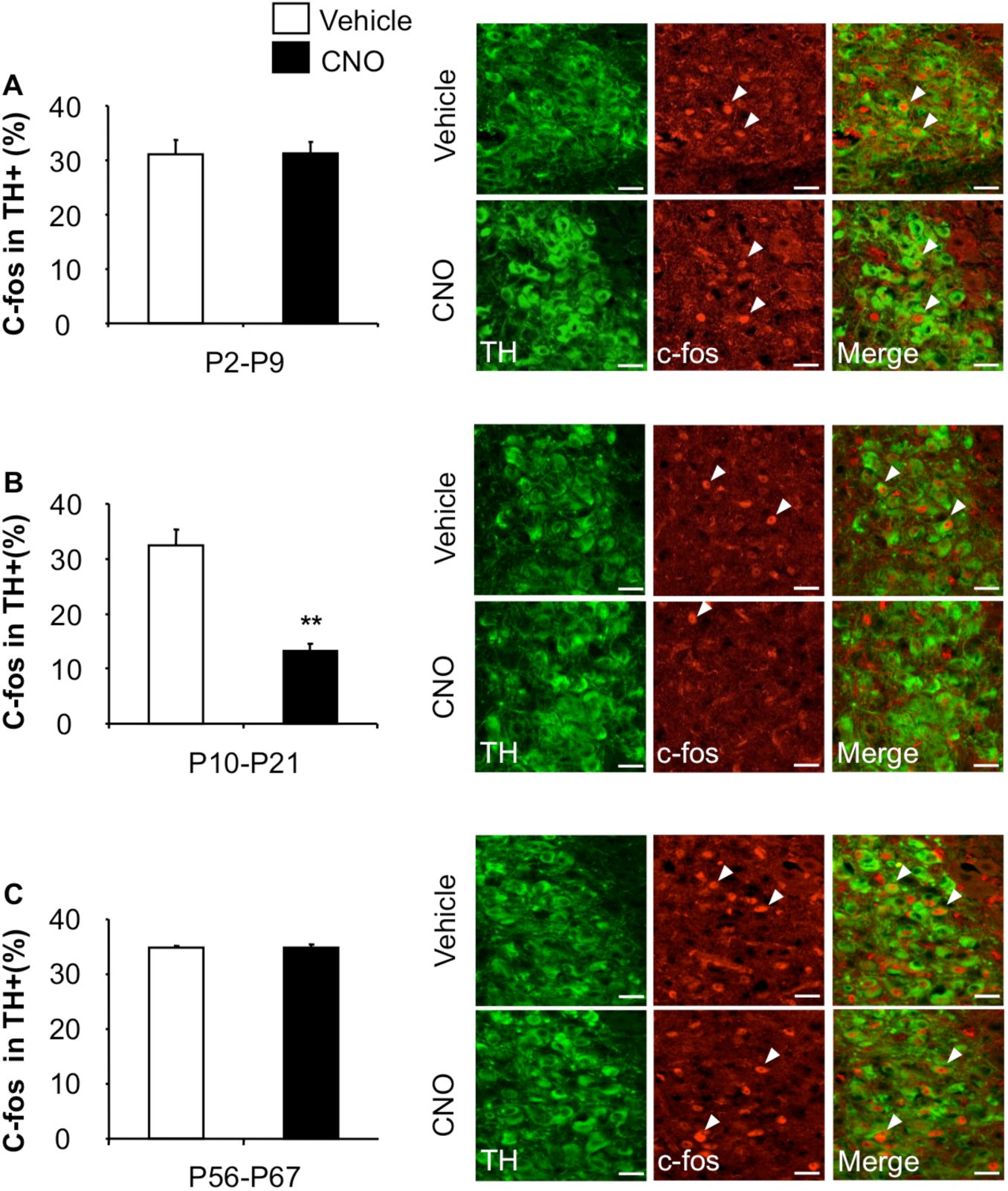
Pharmacogenetic inhibition of NE neurons between P10-21 has long-term consequences for stress-induced LC-NE reactivity. **(A-C)** c-fos and TH double immunostaining in the LC after a forced swim stressor in adult DBH-hM4Di^+^ mice that had previously been treated with CNO and vehicle during different developmental time periods. Single arrows depict single-labeled cells, and double arrows depict double-labeled cells. Scale bars represent 20 μm. Decrease in the percentage of TH+ cells that are also c-fos+ in the P10-P21 **(B)** but not other groups **(A, C)** (one-way ANOVA for main effect of treatment: P2-P9: F_1,8_=0.006, p=0.9395; P10-P21: F_1,7_=37.119, p<0.01; P56-P67: F_1,8_=0.058, p=0.8171; n=4/group). See also Figure S3.

### Inhibition of NE neurons between P10-P21 interferes with α2-A mediated hypothermia in adults

α2-A receptors are the predominant autoreceptors on LC-NE neurons that provide negative feedback to the LC and common heteroreceptors that mediate responses to secreted NE (Langer, 2015). To assess α2-A receptor function in adulthood, mice were acutely treated with the α2-A receptor agonist clonidine, and their core body temperature was measured. Interestingly, only mice treated with CNO during P10-P21, displayed an attenuated hypothermic response when compared to their controls (Supplementary Figure 4A-C). Next, to ensure that the observed changes in thermoregulation were not the result of adaptation in the serotonin system we systemically injected the 5-HT_1A_ agonist 8-OH-DPAT, as 5-HT_1A_ agonist are well known to generate hypothermic responses (Ray et al., 2011). We observed an acute hypothermic response and no differences seen between CNO and vehicle treated groups (Supplementary Figure 4D). These results indicate that interfering with NE activity P10-P21, but not during other periods (P2-P9, P56-P67) results in long-term changes in the sensitivity of α2-A receptors.

### Inhibition of NE neurons between P10-P21 leads to long-term molecular adaptations in the adult LC-NE circuit

Identification of physiological changes in response to an α2-A agonist in adults from only the P10-P21 cohort suggested long-term adaptations within the NE circuitry. We therefore decided to examine the expression of genes critical to NE signaling and regulation in this developmental group and in the reference adult cohort. We found that mRNA expression of LC α2-A (ADRA2A), DBH and NET were all decreased in the P10-P21 CNO-treated animals relative to their vehicle controls. No such changes were seen in the P56-P67 group (Figure 4A-C). Further, we detected no difference in either TH or MAO A mRNA expression in the LC in either cohort (Supplementary Figure 5G, H).

**Figure 4.**
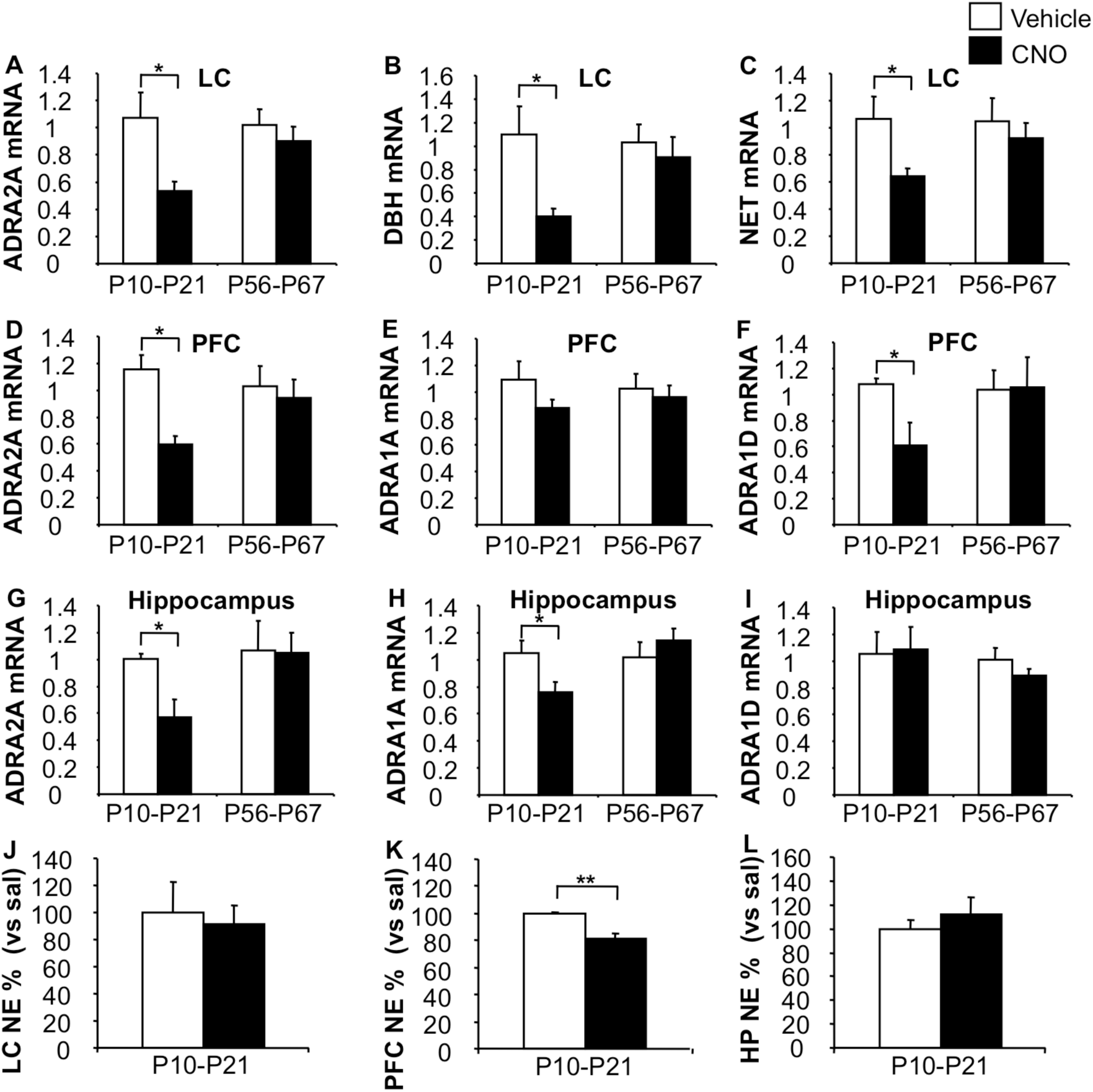
Selective LC-NE system adaptations after pharmacogenetic inhibition of NE neurons. **(A-C)** CNO treatment of DBH-hM4Di^+^ mice between P10-21, but not P56-P67, results in decreased LC (A) ADRA2A (one-way ANOVA for main effect of treatment: P10-P21: F_1,9_=8.108, p<0.05; P56-P67: F_1,7_=0.561, p=0.472; n=5-6/group) (B) DBH (one-way ANOVA for main effect of treatment: P10-P21: F_1,9_=9.112, p<0.05; P56-P67: F_1,7_=0.308, p=0.5963) and (C) NET mRNA expression in the LC (one-way ANOVA for main effect of treatment: P10-P21 F_1,9_=6.624, p<0.05; P56-P67: F_1,7_=0.371, p=0.5619; n=5-6/group). **(D-F)** CNO treatment of DBH-hM4Di^+^ mice between P10-21, but not P56-P67, results in decreased (D) PFC ADRA2A (one-way ANOVA for main effect of treatment: P10-P21: F_1,9_=8.257, p<0.05; P56-P67: F_1,7_=0.170, p=0.6926; n=5-6/group) and (F) PFC ADRA1D mRNA levels (one-way ANOVA for main effect of treatment: P10-P21: F_1,9_=5.569, p<0.05; P56-P67: F_1,7_=0.005, p=0.9482; n=5-6/group), along with no differences in (E) PFC ADRA1A mRNA levels in the adulthood (one-way ANOVA for main effect of treatment F_1,9_=2.378, p=0.1574; P56-P67: F_1,7_=2.583, p=0.1521; n=5-6/group). **(G-I)** CNO treatment of DBH-hM4Di^+^ mice between P10-21, but not P56-P67, results in decreased (G) hippocampal ADRA2A (one-way ANOVA for main effect of treatment P10-P21: F_1,9_=8.686, p<0.05; P56-P67: F_1,7_=0.006, p=0.942; n=5-6/group) and (H) ADRA1A (one-way ANOVA for main effect of treatment P10-P21: F_1,9_=5.957, p<0.05; P56-P67: F_1,7_=0.747, p=0.416, n=5-6/group) along with no changes in ADR1D mRNA levels in adulthood (one-way ANOVA for main effect of treatment P10-P21: F_1,9_=0.014, p=0.9075; P56-P67: F_1,7_=1.494, p=0.2611; n=5-6/group). **(J)** No differences were detected in NE levels in the LC in mice treated with CNO during P10-P21 when comparing them to vehicle controls (one-way ANOVA for main effect of treatment: NE: F_1,8_=0.114, p=0.7439; n=4-6/group). **(K)** In contrast, P10-P21 CNO treatment results in decreased NE levels in the PFC in adulthood when compared to vehicle controls (one-way ANOVA for main effect of treatment: NE: F_1,8_=20.405, p<0.01; n=4-6/group). **(L)** No changes were detected in adult NE levels after P10-P21 CNO intervention in the hippocampus (one-way ANOVA for main effect of treatment: NE: F_1,8_=0.518, p=0.4922; n=4-6/per group). Means are represented as ±SEM. (*p<0.05; **p<0.01). See also Supplementary Figure 4.

We next examined expression levels of NE receptors in LC projection areas that receive extensive NE innervation and have been implicated in the behaviors tested in our earlier studies. Specifically, the prefrontal cortex (PFC), hippocampus, hypothalamus and amygdala were selected. We found that interfering with NE neuronal activity during P10-P21, but not during P56-67, resulted in decreased expression of α2-A and α1-D (ADRA1D) in the PFC (Figure 4D-F) and decreased α2-A and α1-A (ADRA1A) receptor mRNA levels in the hippocampus (Figure 4G-I) of adult animals. In contrast, the expression of these receptors appeared unchanged in the hypothalamus and amygdala for both the P10-P21 and the P56-67 cohorts (Supplementary Figure 5A-F). Thus, changes to key components of the NE circuit were evident in select target areas in adult animals that were treated with CNO between P10-P21, but not in animals treated in adulthood.

Having observed altered mRNA expression of NE receptors, we assessed NE levels in the same brain regions in mice that were exposed to the P10-P21 intervention. We micro-dissected the LC, the PFC, the hippocampus, the hypothalamus and the amygdala to measure total NE content in each region by HPLC. Interestingly, NE content was decreased in the PFC of P10-P21 CNO-treated when compared to vehicle-treated mice (Figure 4K). No such changes were observed in the LC, hippocampus, hypothalamus or amygdala (Figure 4J, L and Supplementary Figure 5I, J). In addition, no differences were detected in serotonin or dopamine levels in these brain regions between CNO or vehicle-treated animals (*Data not shown*).

### Administration of the α2-A receptor agonist, guanfacine, during p10-P21, phenocopies the adult behavioral effects of suppressing NE neuron activity at that time

One important developmental milestone in the NE circuit is the rapid rise of firing rates and the establishment of α-2A receptor mediated negative feedback during P10-P21 (Murrin et al., 2007). Given the extensive changes in α2-A receptor expression and function that we observed in the P10-P21 group treated with CNO, we tested whether stimulation of the α2-A receptor during this time would phenocopy our findings. We decided to use Guanfacine, which is an α2-A agonist that is routinely prescribed in children and adolescents to treat attention deficit hyperactivity disorder (ADHD) (Langer, 2015).

Mice were treated with guanfacine (1 mg/kg) from either P10-P21 or P56-P67 as a reference period (Figure 5A). Adult mice treated with guanfacine during P10-P21, but not P56-P67, displayed decreased percent center distance (Figure 5B, C) along with no changes in the total path travelled in the open-field (Supplementary Figure 6A, B). In addition, mice treated with guanfacine from P10-P21, but not P56-P67, showed decreased percentage time in the open arms in the elevated-plus maze (Figure 5D, E), a finding not detected with the pharmacogenetic approach.

**Figure 5.**
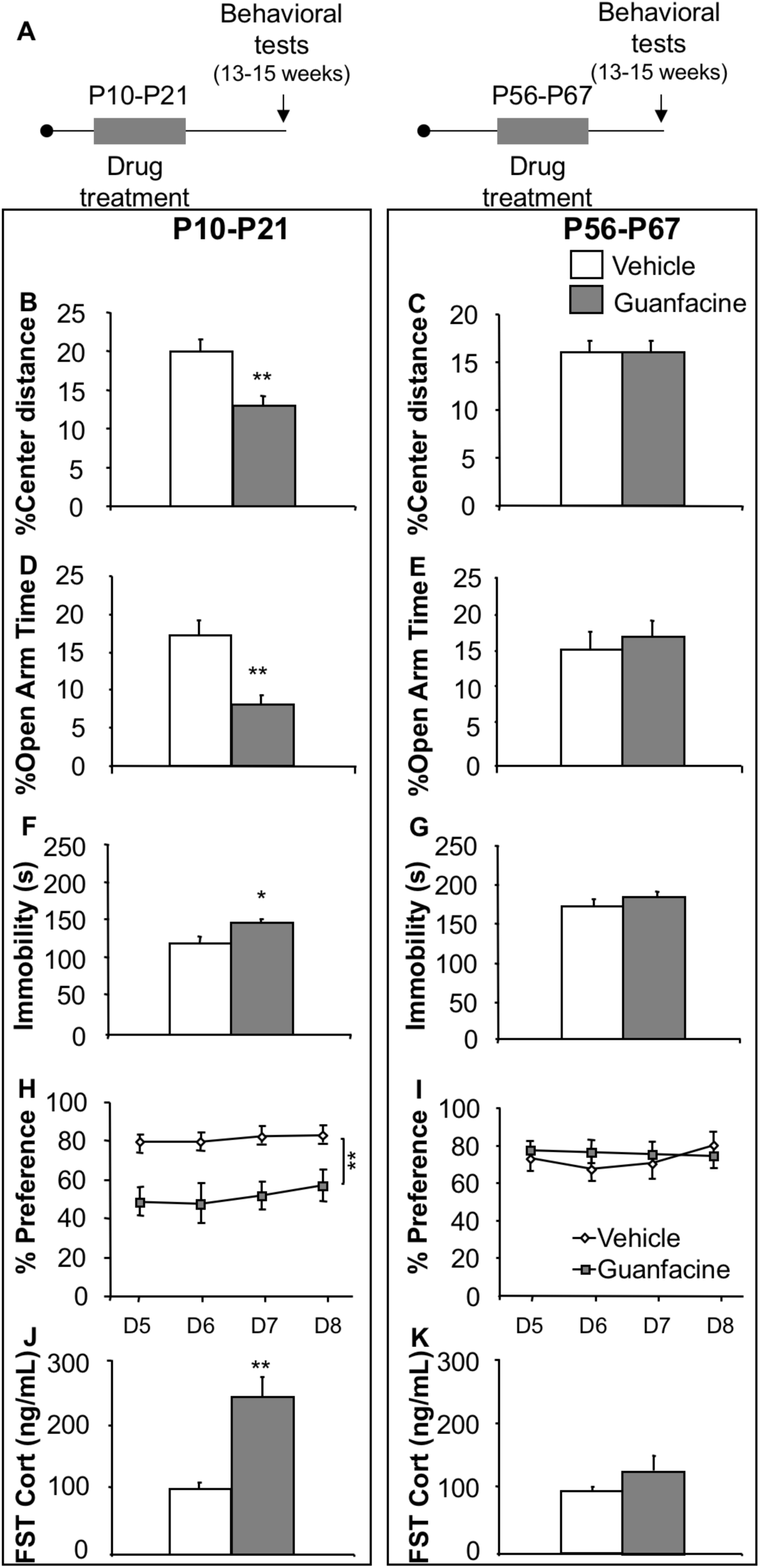
Guanfacine treatment between P10-P21 results in increased anxiety and depression-like behaviors in adulthood. **(A)** Experimental timeline. **(B, C)** P10-P21 guanfacine treatment, but not from P56-P67, results in decreased percent center distance in the open-field test (one-way ANOVA for main effect of treatment P10-P21: F_1,36_=11.072, p<0.01; P56-P67: F_1,28_=0.001, p=0.9742; n=15-20/group), **(D, E)** and decreased percent time in the open arms in the elevated-plus maze test (one-way ANOVA for main effect of treatment P10-P21: F_1,36_=13.146, p<0.01; P56-P67: F_1,28_=0.254, p=0.6184; n=15-20/group). **(F, G)** Increased immobility in the FST in the P10-P21 guanfacine group in adulthood when compared to their control mice. No such difference was observed in the P56-P67 group (one-way ANOVA for main effect of treatment P10-P21: F_1,36_=5.222, p<0.05; P56-P67: F_1,28_=2.015, p=0.1688; n=15-20/group). **(H, I)** P10-P21 guanfacine administration, but not P56-P67, results in decreased sucrose preference (one-way ANOVA repeated measures for main effect of treatment P10-P21: F_1,36_=12.524, p<0.01; P56-P67: F_1,28_=0.033, p=0.8571; n=15-20/group) **(J, K)** Increased post forced swim stress-induced CORT release in P10-P21 guanfacine treated mice, but not in P56-P67, when compared to their respective controls (one-way ANOVA for main effect of treatment P10-P21: F_1,18_=19.660, p<0.01; P56-P67: F_1,17_=0.001, p=0.9742; n=15-20 per group). Means are represented as ±SEM. (*p<0.05; **p<0.01). See also Supplementary Figure 5.

Consistent with our pharmacogenetic approach results testing depression-like behaviors, we also observed decreased mobility in the FST and decrease in sucrose preference in mice that were treated with guanfacine during P10-P21, but not the other time period (Figure 5F-I). Moreover, the P10-P21 group also showed an increased CORT response after a forced-swim stressor (Figure 5J, K). Finally, clonidine induced hypothermia in adulthood was attenuated in P10-P21 mice but not in P56-67 mice (Supplementary Figure 6C, D). In sum, treatment of pups from P10-P21 with the α2-A agonist guanfacine broadly reproduces the effects seen from pharmacogenetically suppressing NE neuronal function during the same developmental time period.

## Discussion

Our results demonstrate the existence of a developmental sensitive period between P10 and P21, during which disruption of NE signaling leads to enduring adult behavioral phenotypes relevant to depression and anxiety. Inhibition of NE signaling using the DREADD system prior to this time (P2-P9) does not result in a long-term behavioral phenotype. Interestingly, this developmental window is characterized by an immature LC that has limited spontaneous activity and a robust and prolonged physiological response to environmental stimuli (Nakamura et al., 1987; Murrin et al., 2007; Raineki et al., 2010). In turn, the P10-P21 period is characterized by emergence of spontaneous activity and a potent α2-A autoreceptor mediated negative feedback (Murrin et al., 2007).

Nevertheless, the LC continues to mature past P21 as NE levels steadily increase until plateauing near adult levels in early adolescence (Murrin et al., 2007). This stabilization of LC function corresponds to our results that by P33 interfering with NE neuronal firing no longer had enduring sequelae. This is further supported by negative results in our P56-P67 cohort. As a result, our data suggest that there is a complex interplay between the emergence of spontaneous NE neuronal activity, the establishment of LC negative feedback and baseline NE levels, which ultimately results in determining adult emotion-like behaviors. The fact that we did not detect behavioral changes in the adolescent and early adulthood interventions, suggests that once mature, the NE system is sufficiently stable to withstand short-term perturbations without eliciting the long-term adaptations seen during P10-P21.

Molecular interrogation of NE machinery in the P10-P21 cohort revealed distinct alterations in the LC and its target regions. The LC was broadly affected including reduced mRNA levels for the rate limiting synthesis enzyme DBH, the reuptake transporter NET, and the inhibitory α2-A autoreceptor responsible for feedback inhibition. In this regard, the absence of change in basal NE levels in the LC raises the possibility that interfering with LC firing during the sensitive period changes LC ability to mount stress responses. Our results indicating that fewer LC TH neurons express c-fos following stress, support this possibility. Furthermore, a change in LC reactivity would be expected to be accompanied by molecular adaptations in NE target regions. Indeed, molecular NE machinery in some LC target regions, such as the PFC, but not others was permanently altered.

LC projects throughout the neocortex and the limbic structures. Remarkably we saw no adaptations in α-adrenergic receptor expression in the amygdala or the hypothalamus. The hypothalamus receives substantial NE innervation from the medullary catecholamine nuclei and the ventral tegmentum rather than solely the LC (Moore and Bloom, 1979; Palkovits et al., 1979; Sawchenko and Swanson, 1982) raising the possibility that the P10-P21 sensitive period is particular to LC NE neurons. This possibility is further supported by our observation that the amygdala, which receives NE inputs from the lateral tegmentum (Moore and Bloom, 1979), did not exhibit molecular changes. In contrast, the hippocampus and the PFC exhibited a robust reduction in α2-A receptor expression. Ascending NE fibers arising from regions other than the LC terminate prior to reaching the hippocampus and the neocortex, which receive their robust innervation from the LC (Moore and Bloom, 1979). Interestingly, only the PFC exhibited a long-lasting decrease in baseline NE suggesting a shift in a homeostatic set point for NE signaling. While it is difficult to infer how different NE levels in the hippocampus and the PFC may translate to behaviors, PFC neuronal activity encodes FST behavioral states and modulates anhedonic behaviors (Warden et al., 2012; Ferenczi et al., 2016). Together the results highlight circuit specific changes that are associated with depression-related but also with some anxiety-related features following inhibition of NE function during a sensitive period.

NE-acting medications are part of the pharmacological arsenal used to treat psychiatric disorders in children. In particular, guanfacine have been used for treating ADHD (Lee, 1997) and clonidine for post-traumatic stress disorder (Harmon and Riggs, 1996). Our studies highlight the critical need to fully understand the implications of treating developing brains with pharmacologic agents targeting modulatory amines. There has been relatively little research on drug by development interactions but given the potential long-term consequences of treatments during sensitive periods, additional consideration by physicians prescribing psychoactive medications to young children seems warranted.

## Funding and disclosure

This work was supported by NIMH R01 MH091844, R56 MH106809 (AD) and NIMH R01 MH91427 (EDL). A Spain Science postdoctoral and Sackler Institute fellowships supported AGG. The authors declare no conflict of interest.

## Acknowledgments

The authors would like to thank Susan Dymecki for her generous gift of the RC∷PDi mice. A.D. and E.D.L. conceived the project. A.G.G., A.D. and E.D.L. wrote the manuscript. Q.M. and A.G.G. performed the experiments.

## References

Armbruster BN, Li X, Pausch MH, Herlitze S, Roth BL (2007) Evolving the lock to fit the key to create a family of G protein-coupled receptors potently activated by an inert ligand. Proc Natl Acad Sci U S A 104:5163–5168.

Ba A, Seri BV (1995) Psychomotor functions in developing rats: ontogenetic approach to structure-function relationships. Neurosci Biobehav Rev 19:413–425.

Carter ME, Yizhar O, Chikahisa S, Nguyen H, Adamantidis A, Nishino S, Deisseroth K, de Lecea L (2010) Tuning arousal with optogenetic modulation of locus coeruleus neurons. Nat Neurosci 13:1526–1533.

Charney DS (2003) Neuroanatomical circuits modulating fear and anxiety behaviors. Acta Psychiatr Scand Suppl:38–50.

Cottingham C, Wang Q (2012) alpha2 adrenergic receptor dysregulation in depressive disorders: implications for the neurobiology of depression and antidepressant therapy. Neurosci Biobehav Rev 36:2214–2225.

Ferenczi EA, Zalocusky KA, Liston C, Grosenick L, Warden MR, Amatya D, Katovich K, Mehta H, Patenaude B, Ramakrishnan C, Kalanithi P, Etkin A, Knutson B, Glover GH, Deisseroth K (2016) Prefrontal cortical regulation of brainwide circuit dynamics and reward-related behavior. Science 351:aac9698.

Garcia-Garcia AL, Meng Q, Richardson-Jones J, Dranovsky A, Leonardo ED (2015) Disruption of 5-HT function in adolescence but not early adulthood leads to sustained increases of anxiety. Neuroscience.

Garcia-Garcia AL, Meng Q, Canetta S, Gardier AM, Guiard BP, Kellendonk C, Dranovsky A, Leonardo ED (2017) Serotonin Signaling through Prefrontal Cortex 5-HT1A Receptors during Adolescence Can Determine Baseline Mood-Related Behaviors. Cell Rep 18:1144–1156.

Geller B, Reising D, Leonard HL, Riddle MA, Walsh BT (1999) Critical review of tricyclic antidepressant use in children and adolescents. J Am Acad Child Adolesc Psychiatry 38:513–516.

Goddard AW, Ball SG, Martinez J, Robinson MJ, Yang CR, Russell JM, Shekhar A (2010) Current perspectives of the roles of the central norepinephrine system in anxiety and depression. Depress Anxiety 27:339–350.

Gong S, Doughty M, Harbaugh CR, Cummins A, Hatten ME, Heintz N, Gerfen CR (2007) Targeting Cre recombinase to specific neuron populations with bacterial artificial chromosome constructs. J Neurosci 27:9817–9823.

Happe HK, Coulter CL, Gerety ME, Sanders JD, O’Rourke M, Bylund DB, Murrin LC (2004) Alpha-2 adrenergic receptor development in rat CNS: an autoradiographic study. Neuroscience 123:167–178.

Harmon RJ, Riggs PD (1996) Clonidine for posttraumatic stress disorder in preschool children. J Am Acad Child Adolesc Psychiatry 35:1247–1249.

Hazell P, O’Connell D, Heathcote D, Henry D (2000) Tricyclic drugs for depression in children and adolescents. Cochrane Database Syst Rev:CD002317.

Klimek V, Stockmeier C, Overholser J, Meltzer HY, Kalka S, Dilley G, Ordway GA (1997) Reduced levels of norepinephrine transporters in the locus coeruleus in major depression. J Neurosci 17:8451–8458.

Konkol RJ, Bendeich EG, Breese GR (1978) A biochemical and morphological study of the altered growth pattern of central catecholamine neurons following 6-hydroxydopamine. Brain Res 140:125–135.

Lahdesmaki J, Sallinen J, MacDonald E, Kobilka BK, Fagerholm V, Scheinin M (2002) Behavioral and neurochemical characterization of alpha(2A)-adrenergic receptor knockout mice. Neuroscience 113:289–299.

Langer SZ (2015) alpha2-Adrenoceptors in the treatment of major neuropsychiatric disorders. Trends Pharmacol Sci 36:196–202.

Lee BJ (1997) Clinical experience with guanfacine in 2-and 3-year-old children with attention deficit hyperactivity disorder. Inf Mental Hlth J 18:300–305.

Loizou LA, Salt P (1970) Regional changes in monoamines of the rat brain during postnatal development. Brain Res 20:467–470.

Madden CJ, Tupone D, Cano G, Morrison SF (2013) alpha2 Adrenergic receptormediated inhibition of thermogenesis. J Neurosci 33:2017–2028.

Moore RY, Bloom FE (1979) Central catecholamine neuron systems: anatomy and physiology of the norepinephrine and epinephrine systems. Annu Rev Neurosci 2:113–168.

Morris MJ, Dausse JP, Devynck MA, Meyer P (1980) Ontogeny of alpha 1 and alpha 2-adrenoceptors in rat brain. Brain Res 190:268–271.

Murrin LC, Sanders JD, Bylund DB (2007) Comparison of the maturation of the adrenergic and serotonergic neurotransmitter systems in the brain: implications for differential drug effects on juveniles and adults. Biochem Pharmacol 73:1225–1236.

Nakamura S, Kimura F, Sakaguchi T (1987) Postnatal development of electrical activity in the locus ceruleus. J Neurophysiol 58:510–524.

Olson VG, Rockett HR, Reh RK, Redila VA, Tran PM, Venkov HA, Defino MC, Hague C, Peskind ER, Szot P, Raskind MA (2011) The role of norepinephrine in differential response to stress in an animal model of posttraumatic stress disorder. Biol Psychiatry 70:441–448.

Palkovits M, Zaborszky L, Brownstein MJ, Fekete MI, Herman JP, Kanyicska B (1979) Distribution of norepinephrine and dopamine in cerebral cortical areas of the rat. Brain Res Bull 4:593–601.

Raineki C, Pickenhagen A, Roth TL, Babstock DM, McLean JH, Harley CW, Lucion AB, Sullivan RM (2010) The neurobiology of infant maternal odor learning. Braz J Med Biol Res 43:914–919.

Rangel S, Leon M (1995) Early odor preference training increases olfactory bulb norepinephrine. Brain Res Dev Brain Res 85:187–191.

Ray RS, Corcoran AE, Brust RD, Kim JC, Richerson GB, Nattie E, Dymecki SM (2011) Impaired respiratory and body temperature control upon acute serotonergic neuron inhibition. Science 333:637–642.

Richardson-Jones JW, Craige CP, Nguyen TH, Kung HF, Gardier AM, Dranovsky A, David DJ, Guiard BP, Beck SG, Hen R, Leonardo ED (2011) Serotonin-1A autoreceptors are necessary and sufficient for the normal formation of circuits underlying innate anxiety. J Neurosci 31:6008–6018.

Sara SJ (2009) The locus coeruleus and noradrenergic modulation of cognition. Nat Rev Neurosci 10:211–223.

Sawchenko PE, Swanson LW (1982) The organization of noradrenergic pathways from the brainstem to the paraventricular and supraoptic nuclei in the rat. Brain Res 257:275–325.

Schramm NL, McDonald MP, Limbird LE (2001) The alpha(2a)-adrenergic receptor plays a protective role in mouse behavioral models of depression and anxiety. J Neurosci 21:4875–4882.

Schwarz LA, Luo L (2015) Organization of the locus coeruleus-norepinephrine system. Curr Biol 25:R1051–1056.

Shishkina GT, Kalinina TS, Sournina NY, Saharov DG, Kobzev VF, Dygalo NN (2002) Effects of antisense oligodeoxynucleotide to the alpha2A-adrenoceptors on the plasma corticosterone level and on elevated plusmaze behavior in rats. Psychoneuroendocrinology 27:593–601.

Snyder K, Wang WW, Han R, McFadden K, Valentino RJ (2012) Corticotropin-releasing factor in the norepinephrine nucleus, locus coeruleus, facilitates behavioral flexibility. Neuropsychopharmacology 37:520–530.

Treadway MT, Zald DH (2011) Reconsidering anhedonia in depression: lessons from translational neuroscience. Neuroscience and biobehavioral reviews 35:537–555.

Valentino RJ, Van Bockstaele E (2008) Convergent regulation of locus coeruleus activity as an adaptive response to stress. Eur J Pharmacol 583:194–203.

Warden MR, Selimbeyoglu A, Mirzabekov JJ, Lo M, Thompson KR, Kim SY, Adhikari A, Tye KM, Frank LM, Deisseroth K (2012) A prefrontal cortex-brainstem neuronal projection that controls response to behavioural challenge. Nature 492:428–432.

