## Supplementary Materials for "Inhibition of norepinephrine signaling during a sensitive period disrupts locus coeruleus circuitry and emotional behaviors in adulthood"

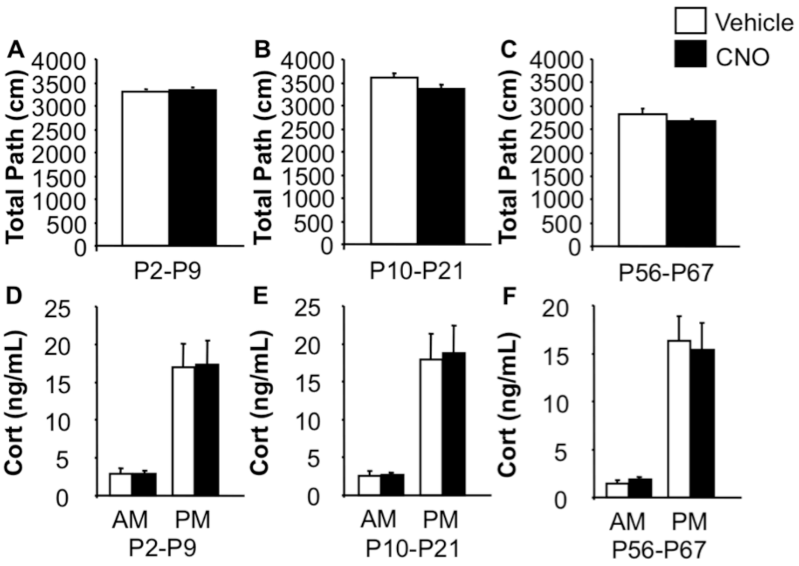


***Supplementary Figure 1. Related to Figure 2. Pharmacogenetic inhibition of NE signaling during development does not impact locomotor activity in adult mice.*** **(A-C)** No changes were detected in total path travelled in open-field in adult mice after CNO treatment during different time windows (one-way ANOVA for main effect of treatment P2-P9: $F_{1,39}$=0.136, p=0.7147; P10-P21: $F_{1,36}$=2.889, p=0.0978; P56-P67: $F_{1,31}$=1.537, p=0.2243; n=14-22/group). **(D-F)** No significant difference between groups in corticosterone levels at the onset of both the light (AM) and the dark phase (PM) (one-way ANOVA P2-P9: AM: $F_{1,16}$=0.001, p=0.9737; PM: $F_{1,16}$=0.004, p=0.9488; P10-P21: AM: $F_{1,16}$=0.024, p=0.8788; PM: $F_{1,16}$=0.024, p=0.8788; P56-P67: AM: $F_{1,16}$=0.028, p=0.8702; PM: $F_{1,16}$=1.120, p=0.3057; n=9-10/group). Means are represented as ±SEM.

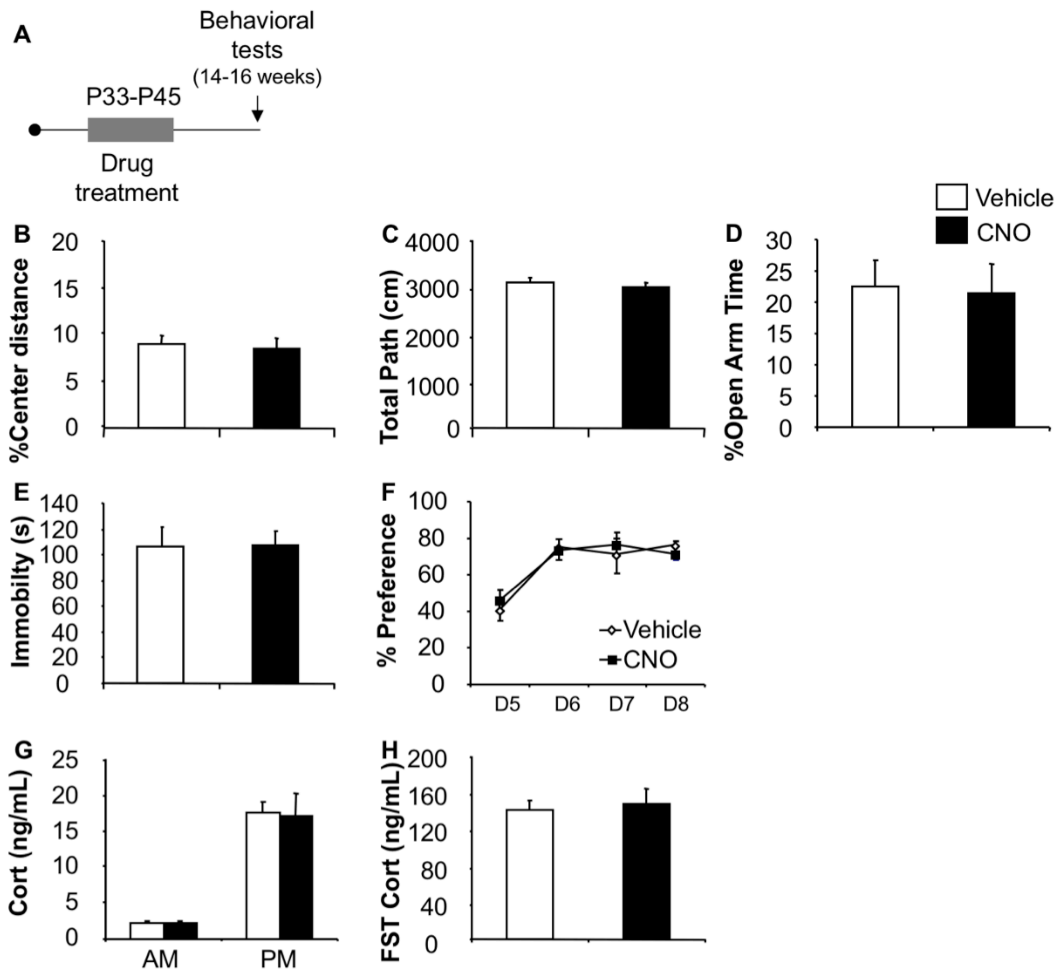


***Supplementary Figure 2. Related to Figure1. Pharmacogenetic inhibition of NE signaling during P33-P45 does anxiety-, depression-related behaviors or stress-induced corticosterone in adult mice.*** **(A)** Experimental timeline. (**B-C)** No difference in percent center distance or total distance travel in the open-field after P33-P45 intervention (one-way ANOVA for main effect of treatment: %center distance: $F_{1,28}$=0.074, p=0.7879; total path: $F_{1,28}$=0.316, p=0.5785; n=15/group). **(D)** No differences were detected in time spent in the open arms in the elevated-plus maze (one-way ANOVA for main effect of treatment: $F_{1,28}$=0.032, p=0.8593; n=15/group). **(E)** Immobility in the FST. No differences were observed in when comparing CNO to vehicle mice (one-way ANOVA for main effect of treatment: $F_{1,30}$=0.030, p=0.869; n=15-17/group). **(F)** Sucrose Preference test. Differences in preference were observed when comparing CNO to vehicle treatment during testing days (Day 5-8) (one-way ANOVA repeated measures for main effect of treatment: $F_{1,26}$=0.013, p=0.9114; n=14/group). **(G)** No significant difference between groups in corticosterone levels at the onset of both the light (AM) and the dark phase (PM) (one-way ANOVA for main effect of treatment: AM:

$F_{1,18}$=0.017, p=0.8981; PM: $F_{1,18}$=0.002; n=9-11/group) **(H)** No changes in forced swim-stress induced CORT (one-way ANOVA for main effect of treatment: P2-P9: $F_{1,18}$=0.156, p=0.6985; n=9-11/group). Means are represented as ±SEM.

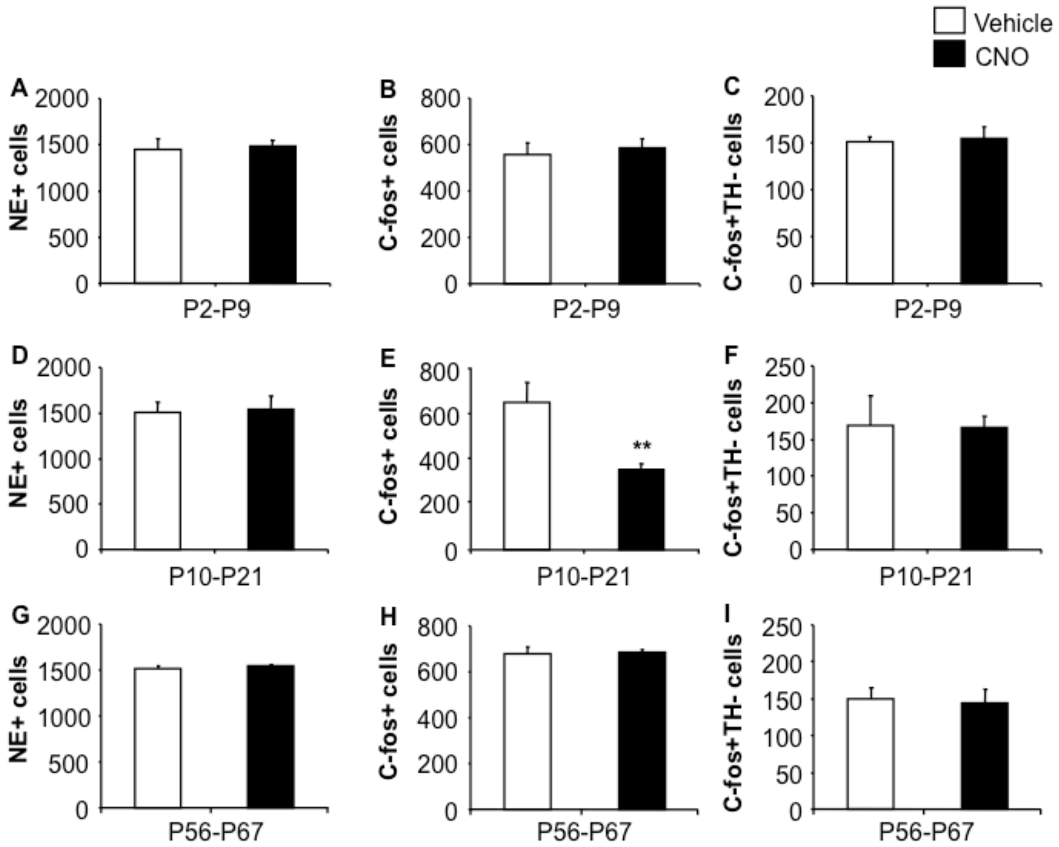


***Supplementary Figure 3. Related to Figure 3. Pharmacogenetic inhibition of NE neurons between P10-P21 alters the LC response to stress in adulthood.*** **(A)** No changes were observed in the number of NE (TH+) neurons (one-way ANOVA for main effect of treatment: $F_{1,8}$=0.050, p=0.8293), **(B)** stress induced c-fos+ (one-way ANOVA for main effect of treatment: $F_{1,8}$=0.219, p=0.6520) and **(C)** c-fos+TH- (one-way ANOVA for main effect of treatment: $F_{1,8}$=0.075, p=0.7907) cells in the LC of adult mice after P2-P9 NE signaling inhibition. **(D)** P10-P21 NE inhibition does not affect the number of NE (TH+) neurons (one-way ANOVA for main effect of treatment: $F_{1,7}$=0.028, p=0.8710), **(E)** but results in a decreased number of stress induced c-fos+ cells (one-way ANOVA for main effect of treatment: $F_{1,7}$=13.416, p<0.01), **(F)** along with no changes in the number of c-fos+TH- cells (one-way ANOVA for main effect of treatment: $F_{1,7}$=0.010, p=0.9243). **(G)** No changes were observed in the number of NE (TH+) (one-way ANOVA for main effect of treatment: $F_{1,8}$=0.526, p=0.4956), **(H)** stress induced c-fos+ (one-way ANOVA for main effect of treatment: $F_{1,8}$=0.018, p=0.8984) and **(I)** c-fos+TH- (one-way ANOVA for main effect of treatment: $F_{1,8}$=0.065, p=0.8070) cells in the LC of adult mice after P56-P67 CNO treatment of DBH-hM4Di$^{+}$ mice (n=4/group for all measures). Means are represented as ±SEM. (*p<0.05; **p<0.01).

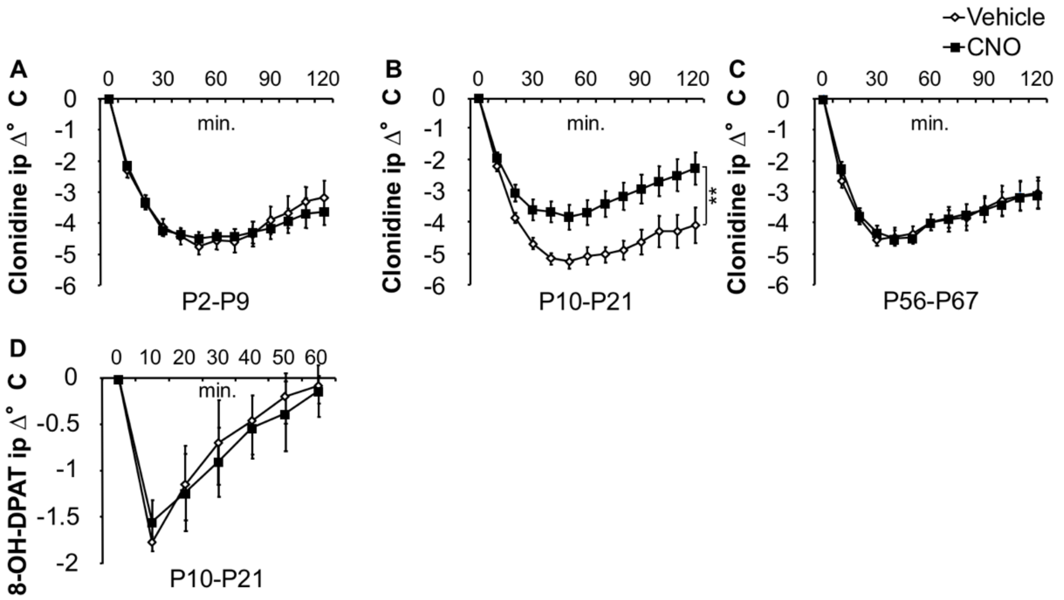


***Supplementary Figure 4. LC-NE system mediated hypothermic responses in adult mice.*** **(A-C)** Hypothermic response to clonidine in adult DBH-hM4Di+ mice treated with CNO and vehicle during different developmental time periods. Decreased hypothermic response to clonidine in P10-P21 (B) but no other groups (A, C) (one-way ANOVA repeated measures for main effect of treatment: P2-P9: $F_{1,16}$=0.017 p=0.8976; P10-P21: $F_{1,17}$=9.227 p<0.01; P56-P67: $F_{1,18}$=0.007 p=0.9349; n=9-11/group). **(D)** No changes were observed in 8-OH-DPAT (5-$HT_{1A}$ agonist) induced hypothermia in the P10-P21 group (one-way ANOVA repeated measures for main effect of treatment: $F_{1,8}$=0.179 p=0.6838; n=5/group). Means are represented as ±SEM. *p<0.05; **p<0.01).

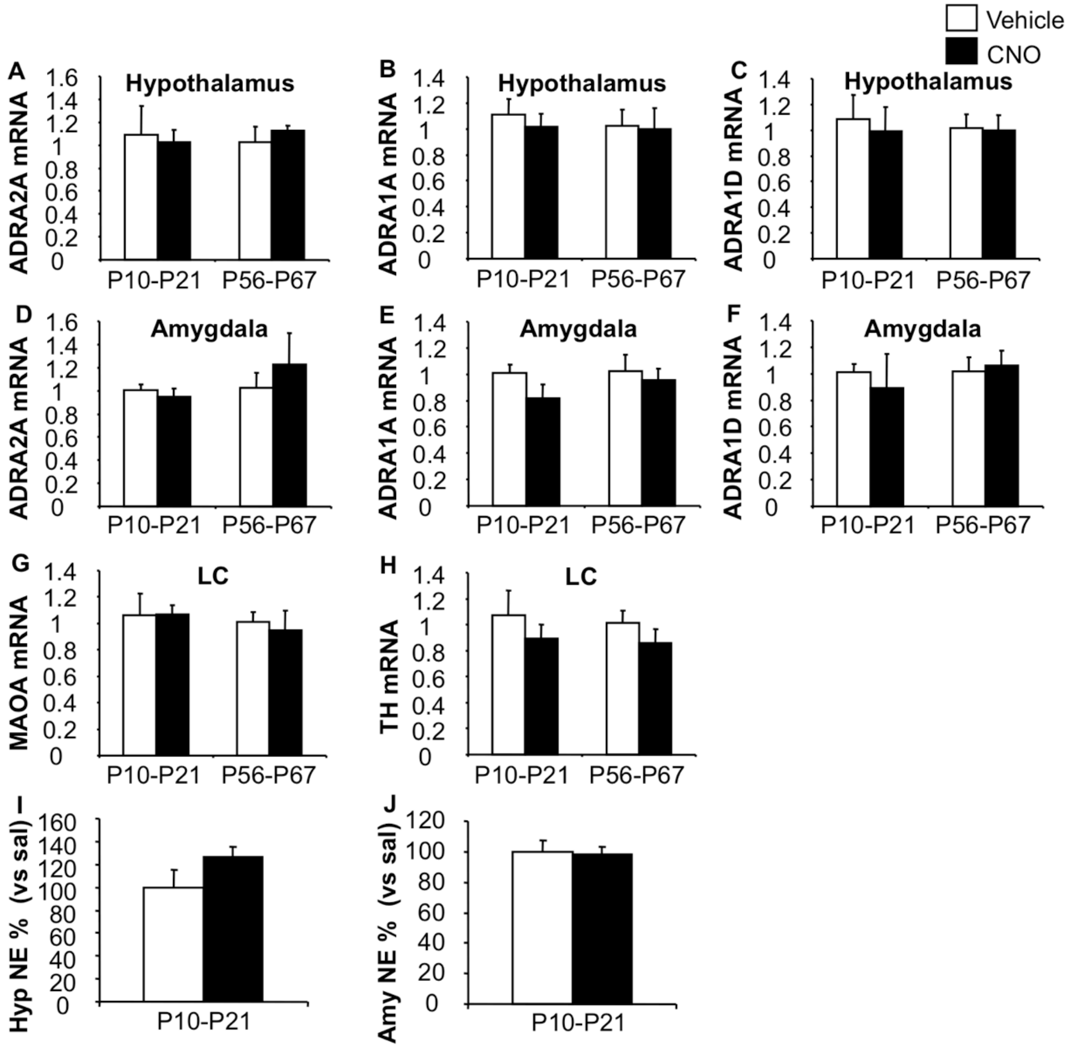


***Supplementary Figure 5. Related to Figure 4.*** **Lack of LC-NE system adaptations after pharmacogenetic inhibition of NE neurons. (A-C)** No changes were observed in the hypothalamus after P10-P21 or P56-P67 intervention in ADRA2A (one-way ANOVA for main effect of treatment: P10-P21: $F_{1,9}$=0.069, p=0.7988; P56-P67: $F_{1,7}$=0.517, p=0.4955), ADRA1A (one-way ANOVA for main effect of treatment: P10-P21: $F_{1,9}$=0.344, p=0.5719; P56-P67: $F_{1,7}$=0.014, p=0.9093) or ADRA1D mRNA levels (one-way ANOVA for main effect of treatment: P10-P21: $F_{1,9}$=0.122, p=0.7351; P56-P67: $F_{1,7}$=0.015, p=0.9046). **(D-F)** No changes were observed in the amygdala after P10-P21 or P56-P67 intervention in ADRA2A (one-way ANOVA for main effect of treatment: P10-P21: $F_{1,9}$=0.373, p=0.5567; P56-P67: $F_{1,7}$=0.380, p=0.5571), ADRA1A (one-way ANOVA for main effect of treatment: P10-P21: $F_{1,9}$=2.135, p=0.1780; P56-P67: $F_{1,7}$=0.191, p=0.6754) or ADRA1D mRNA levels (one-way ANOVA for main effect of treatment P10-P21: $F_{1,9}$=0.168, p=0.6912; P56-P67: $F_{1,7}$=0.069, p=0.8004). **(G)** In the LC, no changes were observed in adult

MAO A (one-way ANOVA for main effect of treatment: P10-P21: $F_{1,9}$=0.006, p=0.9397; P56-P67: $F_{1,7}$=0.108, p=0.7524) or **(H)** TH (Tyrosine hydroxylase) (one-way ANOVA for main effect of treatment P10-P21: $F_{1,9}$=0.723, p=0.4171; P56-P67: $F_{1,7}$=1.164, p=0.3164) mRNA levels after P10-P21 and P56-P67 NE signaling inhibition. **(I)** No changes in NE levels were observed in the hypothalamus (hyp) (one-way ANOVA for main effect of treatment: $F_{1,8}$=2.141, p=0.1815) or **(J)** in the amygdala (amy) (one-way ANOVA for main effect of treatment: NE: $F_{1,8}$=0.041, p=0.8442). (n=4-6/group for all measures). Means are represented as ±SEM. (*p<0.05; **p<0.01).

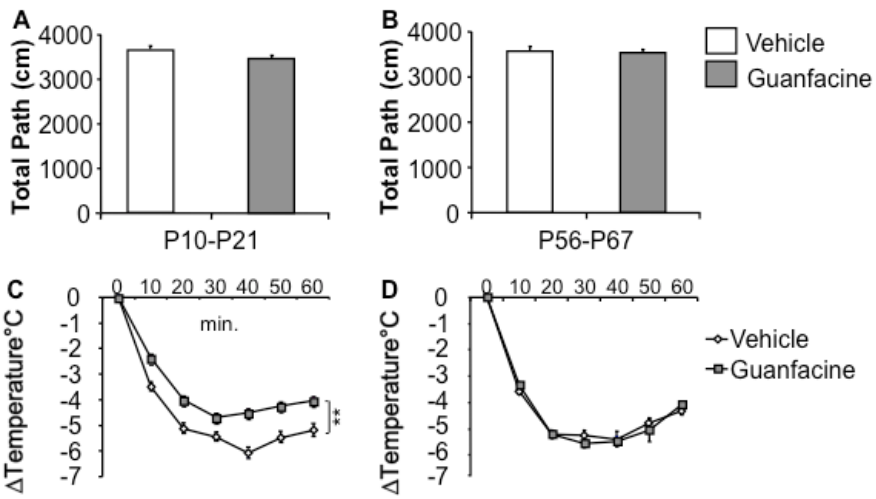


***Supplementary Figure 6. Related to Figure 5. Guanfacine administration during P10-P21 or P56-P67 developmental time windows does not impact locomotor activity in adult mice.*** No changes were detected in total path travelled in open-field in adult mice that were treated with guanfacine during **(A)** P10-P21 (one-way ANOVA for main effect of treatment: P10-P21: $F_{1,36}$=2.479, p=0.1242) and **(B)** P56-P67 (one-way ANOVA for main effect of treatment: P56-P67: $F_{1,28}$=0.301, p=0.5878). **(C, D)** Guanfacine from P10 to P21, but not from P56 to P67, leads to a decreased hypothermic response to Clonidine in adulthood (one-way ANOVA repeated measures for main effect of treatment: P10-P21: $F_{1,8}$=124.058, p<0.01; P56-P67: $F_{1,8}$=0.066, p=0.8041). (n=15/group for all measures). Means are represented as ±SEM. (*p<0.05; **p<0.01).

**SUPPLEMENTARY METHODS AND MATERIALS**

*Animal husbandry*

Animals were housed in groups of three to five per cage and had ad libitum access to food and water. Animals were maintained on a 12:12 light/dark schedule; all testing was conducted during the light period. Animal protocols were approved by the Institutional Animal Care and Use Committee and were conducted in accordance to the NIH Guide for the Care and Use of Laboratory Animals. Care was taken to minimize the number of animals used and their suffering.

*Drugs treatment and administration*

Clozapine-N-oxide (CNO) was obtained from the NIH as part of the Rapid Access to Investigative Drug Program funded by the NINDS. It was dissolved in 1% DMSO and 0.9% Saline.

For hypothermia in figure 1 CNO was injected at a dose of 5 mg/kg in vehicle (0.9% Saline with 1% DMSO) intraperitoneally (i.p.) in adulthood.

For developmental interventions, DBH-hM4Di+ male mice were treated daily with vehicle or CNO i.p. (5 mg/kg in 1% DMSO and 0.9% saline) during different time windows (P2–P9/ P10-P21/ P33-P45/ P56-P67). Specifically, 10 μL per 1 gr of mouse body weight was injected from a 0.5 mg/mL stock solution. For preweaning treatments (P21), the entire litters were removed from dams and placed in a small tray containing bedding from the respective home cage. The tray placed on a scale allow us to measure the weight of an individual pup when removing it for injection. Mice were injected in a random order and immediately placed back in the home cage. Because we aimed at keeping a minimal interference within the litters we assigned all mice within a litter to the same treatment. All mice were housed in the same room and rack and were weaned on P21 and housed in groups of five mice.

For clonidine-induced hypothermia (Supplementary Figure 4), clonidine (0.5 mg/kg in 0.9% saline)(Sigma–Aldrich, St. Louis, MO, USA) was administered i.p. For 8-OH-DPAT induced hypothermia, 1mg/kg in 0.9% saline (Sigma–Aldrich, St. Louis, MO, USA) was injected i.p. 8-OH-DPAT induced hypothermia in mice is dependent on functional 5-$HT_{1A}$ autoreceptors (2).

Guanfacine was dissolved in vehicle (0.9% saline) and administered daily (3-5pm) at a dose of 1 mg/kg i.p. Treatment assignment was randomized in each cage.

*Behavioral and physiological studies*

All animals used for behavioral testing were age matched within 2 weeks. Animals were initially tested at 13–15 weeks of age in different behavioral paradigms in the following order: open-field, elevated-plus maze, sucrose preference and forced swim test and hypothermia with a minimum of 2-3 days between each test. All behavioral testing took place during the light cycle.

*Immunohistochemistry*

*Tissue processing*

After anesthesia with ketamine and xylazine (100 mg/ml ketamine; 20 mg/ml xylazine), mice were perfused transcardially (cold 0.1 M phosphate buffer (pH = 7.4)(PBS) followed by 4% paraformaldehyde (PFA). Brains were removed, post-fixed (24 h), cryoprotected in a 30% sucrose solution (in phosphate buffer) and stored at 4 °C. Serial sections (35 μM) were cut through the entire brain on cryostat (Leica CM3050 S) and stored in PBS with 0.1% NaN3. Specifically, sections at 1:6 interval trough the LC were used.

*Immunofluorescence studies*

For visualization purposes, free floating coronal serial sections (35 μm) of the LC were first washed with PBS 3x10 min followed by 30 min incubation with Triton 1% in PBS. Afterwards, sections were blocked in 10% NDS for 1 hour at room temperature and incubated in primary antibodies for 72 hrs at 4°C (1:100, rabbit anti-HA (Invitrogen, Camarillo, CA), 1:1000, sheep anti-tyrosine hydroxylase (TH) (Abcam)). After washing with PBS, sections were incubated for 1 hr with the secondary antibody donkey anti-rabbit biotin (1:200)(Jackson ImmunoResearch, West Grove, PA)) followed by amplification with avydin (1:200, Cy3 (Jackson ImmunoResearch, West Grove, PA)) complex and donkey anti-sheep cy2 (1:200) (Jackson ImmunoResearch, West Grove, PA) and NeuroTrace fluorescent Nissl stain (Invitrogen, Grand Island, NY). Thereafter, sections were washed 2x10min followed by a last 30 min. wash and mounted on glass slides and embedded with Prolong Gold Antifade Reagent (Invitrogen, Grand Island, NY).

For the stress-induced c-fos experiments, c-fos was induced as previously described by a forced swim stressor (10 min) and mice were perfused 2 hours after stress (6). After tissue processing, LC sections were washed with PBS 3x10min followed by incubation with Triton 1% in PBS for 30 min. Afterwards, serial sections were blocked in 10% Normal donkey serum (NDS) and 1% Triton in PBS for 1 hr. and then incubated with rabbit c-fos antibody (1:5000, Millipore) and sheep TH (1:1000, Abcam) overnight at 4 °C. After washing with PBS, sections were incubated for 1 hr with secondary antibody (1:200 donkey anti-rabbit biotin (Jackson ImmunoResearch, West Grove, PA)) followed by amplification with avidin (1:200, Cy3 (Jackson ImmunoResearch, West Grove, PA)) complex and donkey anti-sheep (1:200, Cy2 (Jackson ImmunoResearch, West Grove, PA). Thereafter, sections were washed 2x10min followed by a last 30 min. wash and mounted on glass slides and embedded with Prolong Gold Antifade Reagent (Invitrogen, Grand Island, NY).

*Image processing and quantification*

For all experiments, sections were imaged with identical exposure times, and parameters with a confocal microscope (Leica, NY, USA) at a magnification of 20x. 4-5 LC sections per mouse were counted. Each section was assessed for the number of single c-fos+ and TH+ cells as well as the number of double-labeled cells. Co-localization of induced c-fos immunoreactivity with TH cell bodies was confirmed with a stack analysis of the images and evaluating the sections in Z series (ImageJ software).

*Quantitative PCR*

Total RNA from the tissues was extracted using TRIzol (Life Technologies, Grand Island, NY, USA). The SuperScript® III First-Strand Synthesis System (Life Technologies, Grand Island, NY, USA) was used to synthesize cDNA, and PCR was performed and quantified using SYBR Green real-time PCR Master Mix (Life Technologies, Grand Island, NY, USA). Analysis was performed for Adrenergicα2A receptor (ADRA2A), Adrenergicα1A receptor (ADRA1A), Adrenergicα1D receptor (ADRA1D), monoamine oxidase A (MAOA) and TH mRNA expression. The β-actin mRNA expression was analyzed as the internal control. Primers used in the real-time quantitative PCR were shown as follows. For ADRA2A, the sense primer was 5'-CTGGACACGGACCTGCTT-3' and the antisense primer was 5'-GAGGCTTCATTTCCTTCTGC-3'. For ADRA1A, the sense primer was 5'-TCAATGAGGAGCCAGGATA-3' and the antisense primer was 5'-GATACGGAGCGTCACTTGC-3'. For ADRA1D, the sense primer was 5'-GTGTCCAGCCTGTCCCATAA-3' and the antisense primer was 5'-CGTCTTGGGGAACATTTAGG-3'. For MAOA, the sense primer was 5'-CTCGGATATTCTCAGTCACCA-3' and the antisense primer was 5'-GAGGACCATTATCTGTTCACTTATT-3'. For TH, the sense primer was 5'-GTCTACTGTCTGCCCGTGAT-3' and the antisense primer was 5'-CAATGTCCTGGGAGAACTGG-3'. For GAPDH the sense primer was 5'-GCCTTCCGTGTTCCTACCC-3' and the antisense primer was 5'-

TGAAGTCGCAGGAGACAACC-3’. For β-actin, the sense primer was 5’-GACGGCCAGGTCATCACTAT-3’ and the antisense primer was 5’-ATGCCACAGGATTCCATACC-3’.

*Statistical Analysis*

All statistical analyses were performed using Stat View (SAS Institute Inc.). Final group numbers are shown in Figure legends. Results from data analyses were expressed as mean ± SEM. $p<0.05$ was used as the threshold for significance. Group differences were analyzed using a one-way analysis of variance (ANOVA) unless otherwise stated. Repeated measures ANOVA was used for hypothermia and sucrose preference experiments.

**SUPPLEMENTARY REFERENCES**

1. Ray RS, Corcoran AE, Brust RD, Kim JC, Richerson GB, Nattie E, et al. (2011): Impaired respiratory and body temperature control upon acute serotonergic neuron inhibition. *Science*. 333:637-642.
2. Richardson-Jones JW, Craige CP, Nguyen TH, Kung HF, Gardier AM, Dranovsky A, et al. (2011): Serotonin-1A autoreceptors are necessary and sufficient for the normal formation of circuits underlying innate anxiety. *J Neurosci*. 31:6008-6018.
3. Garcia-Garcia AL, Meng Q, Canetta S, Gardier AM, Guiard BP, Kellendonk C, et al. (2017): Serotonin Signaling through Prefrontal Cortex 5-HT1A Receptors during Adolescence Can Determine Baseline Mood-Related Behaviors. *Cell Rep*. 18:1144-1156.
4. Garcia-Garcia AL, Meng Q, Richardson-Jones J, Dranovsky A, Leonardo ED (2015): Disruption of 5-HT function in adolescence but not early adulthood leads to sustained increases of anxiety. *Neuroscience*.
5. David DJ, Samuels BA, Rainer Q, Wang JW, Marsteller D, Mendez I, et al. (2009): Neurogenesis-dependent and -independent effects of fluoxetine in an animal model of anxiety/depression. *Neuron*. 62:479-493.
6. Garcia-Garcia AL, Venzala E, Elizalde N, Ramirez MJ, Urbiola A, Del Rio J, et al. (2013): Regulation of serotonin (5-HT) function by a VGLUT1 dependent glutamate pathway. *Neuropharmacology*. 70:190-199.
